# Application of B cell immortalization for the isolation of antibodies and B cell clones from vaccine and infection settings

**DOI:** 10.1101/2022.03.29.485179

**Authors:** Kristin L. Boswell, Timothy Watkins, Evan M. Cale, Jakob Samsel, Sarah F. Andrews, David R. Ambrozak, Jefferson I. Driscoll, Michael A. Messina, Sandeep Narpala, Christine S. Hopp, Alberto Cagigi, Joseph P. Casazza, Takuya Yamamoto, Tongqing Zhou, William R. Schief, Peter D. Crompton, Julie E. Ledgerwood, Mark Connors, Lucio Gama, Peter D. Kwong, Adrian McDermott, John R. Mascola, Richard A. Koup

## Abstract

The isolation and characterization of neutralizing antibodies from infection and vaccine settings will inform future vaccine design, and methodologies that streamline the isolation of antibodies and the generation of B cell clones are of great interest. Retroviral transduction to express Bcl-6 and Bcl-xL in primary B cells has been shown to promote long-term B cell survival and antibody secretion *in vitro*, and can be used to isolate antibodies from memory B cells. The application of this methodology to B cell subsets from tissues and to B cells from individuals with chronic infection has not been extensively characterized. Here, we characterize Bcl-6/Bcl-xL B cell immortalization across multiple tissue types and B cell subsets in healthy and HIV-1 infected individuals, as well as individuals recovering from malaria. In HIV-1- and malaria-uninfected donors, naïve and memory B cell subsets from PBMC and tonsil tissue transformed with similar efficiencies, and displayed similar characteristics after transformation with respect to their longevity and immunoglobulin secretion. In HIV-1-viremic individuals or in individuals after malaria infection, the CD27^-^CD21^-^ memory B cell subsets transformed with lower efficiencies compared to the CD27^+^CD21^+^ populations, but following transformation B cells expanded and secreted IgG with similar efficiency. Using B cells from HIV-1-infected individuals, we combined Bcl-6/Bcl-xL B cell immortalization with a HIV-1 microneutralization assay to isolate broadly neutralizing antibodies related to VRC13 and VRC38.01. Overall, Bcl-6/Bcl-xL B cell immortalization can be used to isolate antibodies and generate B cell clones from multiple different B cell populations, albeit with different efficiencies.

## Introduction

The isolation and characterization of antibodies from infection settings and vaccine trials in both humans and animal models can inform rational vaccine development. A better understanding of neutralizing antibodies, including their longitudinal maturation and binding specificity can aid in the development of new antigens and vaccine regimens. In addition, several neutralizing antibodies have either been approved for use, or are currently being tested for use as therapeutics to prevent or treat infectious diseases, such as in the cases of Ebola virus, respiratory syncytial virus (RSV), SARS-CoV-2 and Human Immunodeficiency virus (HIV) (1-4). Thus, efficient strategies to isolate antibodies from memory B cells following infection or vaccination are needed to not only inform future vaccine design, but also to identify novel therapeutics.

Multiple strategies exist to isolate antigen-specific B cells (5). One strategy includes using a recombinant fluorophore-labeled antigen probe to isolate antigen-specific B cells by flow cytometry for subsequent sequencing of the VH region (6, 7). While this strategy enriches for B cells based on antigen binding, antibody function can only be tested after antibody sequencing, cloning, expression and purification. Another method relies on the culture of B cells *in vitro*, where after 10-14 days of culture with cytokines and CD40 Ligand, activated B cells secrete immunoglobulin into culture supernatant and differentiate into plasma cells before cell death (8). This strategy enables screening for antibody function using immunoglobulin-containing culture supernatant, but the small-scale volume of culture supernatant available limits the breadth of screening that can be performed.

An alternative method that can combine both antigen-binding for B cell enrichment and screening of immunoglobulin-containing culture supernatant for antibody function has been applied to isolate rare antibodies from IgG^+^ and IgM^+^ memory B cells in PBMC (9-11). This method relies on the retroviral expression of the transcription factor Bcl-6 and anti-apoptotic molecule Bcl-xL in primary memory B cells. When cultured in the presence of IL-21 and CD40 ligand, memory B cells transformed by Bcl-6 and Bcl-xL expression survive long-term *in vitro*, secrete antibodies into the culture supernatant, and continue to express the B cell receptor (BCR) on the cell surface (9). Thus, Bcl-6/Bcl-xL immortalization of memory B cells provides a flexible tool for antibody isolation where one can enrich for antigen-specific B cells by flow cytometry using a fluorophore-labeled antigen probe, and also screen for antibody function with a renewable source of culture supernatant.

The efficiency and application of Bcl-6 and Bcl-xL expression in specific B cell subsets has not been fully explored, and in select conditions this technology may be limited. For example, in cases where biopsy samples or fine needle aspirates of draining lymph nodes are available, antigen-specific B cell responses from B cell follicles and germinal centers can be studied. Germinal center B cells isolated from secondary lymphoid organs (SLO), however, more readily undergo cell death during *in vitro* culture and antibody discovery techniques that rely on long-term B cell culture may be limited (12). In chronic infection settings such as HIV-1, malaria or tuberculosis, memory B cell populations characterized by the low expression of CD27 and CD21, sometimes referred to as “atypical”, “tissue-like” or “exhausted”, can expand to comprise up to 40-50% of the memory B cell pool in PBMC (13-17). In general, CD27^-^CD21^-^ B cell populations are characterized by the high expression of inhibitory receptors and have an impaired ability to proliferate and produce antibodies when stimulated (13, 18). In the cases of HIV-1 and malaria, antigen-specific B cells can be found within CD27^-^CD21^-^ B cell subsets, and isolation and characterization of antibodies from these subsets is of interest (19, 20). If Bcl6/Bcl-xL induced B cell immortalization can be applied to CD27^-^CD21^-^ B cell populations, this may overcome the lack of B cell proliferation and limited antibody secretion. However, some level of B cell activation and proliferation must occur *in vitro* to enable retroviral transduction and Bcl-6 and Bcl-xL expression. Hence, B cell subsets that have a limited ability to proliferate and/or readily undergo *in vitro* cell death may not be as amenable to B cell transformation.

One example where the application of B cell immortalization in chronic infection is of particular interest, is in the isolation of broadly neutralizing antibodies (bNAbs) from HIV-1-infected individuals. Only 10-25% of HIV-1-infected patients develop cross-reactive neutralizing antibodies, where antibodies found in the sera target conserved regions on the HIV-1 envelope and can neutralize multiple different strains of HIV-1 (21). Neutralization breadth tends to arise in HIV-1-infected individuals with high viral loads after many years of infection (22, 23). If B cells from HIV-1-infected individuals with neutralization breadth can be immortalized, the isolation of highly potent and broadly neutralizing antibodies may be able to be streamlined. B cell immortalization can generate a long-lasting source of B cells that can be screened for binding to multiple different HIV-1 envelope probes, and in addition, larger volumes of culture supernatant allow for screening for neutralization against multiple different HIV-1 pseudoviruses. This would enable a more high-throughput screening of breadth before cloning and sequencing antibodies of interest.

In addition to antibody isolation from memory B cell subsets, immortalized B cells can be used for other applications such as the characterization of naïve B cell repertoires as well as the generation of B cell clones that can serve as experimental controls. In the HIV-1 vaccine field, several novel immunogens have been designed to bind to and expand germline precursor B cells of bNAbs (24, 25). One such germline-targetting immunogen, eOD-GT8, is being examined for its ability to activate rare precursor B cells of the VRC01-class of antibodies, which are CD4-binding site bNAbs characterized by use of the heavy chain gene IGHV1-2 paired with a light chain CDR3 of only 5 amino acids (26, 27). Thus, a rigorous and reproducible methodology to determine not only the frequency of rare VRC01 precursor B cells but also to accurately determine the BCR sequence of these cells of is of vital interest. And while the VRC01 precursor repertoire has been characterized in some subjects in part using a Bcl-6/Bcl-xL-based B cell immortalization methodology (27), the importance of using clonal precursor B cell lines to serve as experimental controls should not be understated. B cell clones can function as controls for testing reagent sensitivity, specificity, determining the level of detection and accounting for operator variability when sorting rare B cells by flow cytometry using B cells probes.

Here, we explore the efficacy of Bcl-6 and Bcl-xL-mediated B cell immortalization to isolate B cell clones and antibodies from naïve and memory B cell subsets obtained from PBMC and tonsil tissue. We characterize the efficiency of retroviral transduction, quantify immunoglobulin secretion, explore the frequency of mutations acquired during *in vitro* culture, and demonstrate that this technology can be implemented to efficiently generate antigen-specific clonal B cell lines from both naïve and memory B cell subsets. In addition, we compare the efficiency of Bcl-6/Bcl-xL B cell immortalization of CD27^+^CD21^+^ and CD27^-^CD21^-^ B cells from HIV-1-infected individuals and from convalescent individuals one week after malaria treatment, and demonstrate a strategy to isolate HIV-1-specific neutralizing antibodies by combining Bcl-6/Bcl-xL B cell immortalization with the TZM-bl neutralization assay. Overall, we found Bcl-6 and Bcl-xL-mediated B cell immortalization to be a valuable tool to isolate antibodies and B cell clones in both vaccine and chronic infection settings.

## Results

### Activation and transduction of multiple B cell subsets from PBMC and tonsil

To better understand the application of Bcl-6 and Bcl-xL B cell transformation to different B cell subsets, we compared the efficiency of Bcl-6/Bcl-xL transformation of naïve and memory B cells isolated from human PBMC, and naïve and memory subsets isolated from human tonsil. For PBMC, we sorted IgD^+^ (naïve) and IgD^-^IgM^-^IgG^+^ (IgG^+^ memory) B cells and activated them in the presence of recombinant human IL-21 (25 ng/ml) and irradiated 3T3-msCD40-ligand (CD40L) feeder cells for 2 days (Figure 1A). We spinoculated (1200*xg*, 32°C, 1 hour) activated B cells with fresh viral supernatant supplemented with polybrene (4 μg/mL) on days 2 and 3 of activation (Supplemental Figure 1). On day 4 post-transduction we observed transduction efficiencies between 45-80% for both IgD^+^ and IgD^-^IgM^-^IgG^+^ B cell populations as determined by the expression of green fluorescent protein (GFP) (Figure 1C, E). For tonsil tissue, we sorted IgD^+^ (naïve) B cells, IgD^-^CD20^+^CD38^+^IgG^+^ (Germinal Center (GC) IgG^+^) B cells, and IgD^-^CD20^+^CD38^-^IgG^+^ (non-germinal center (Non-GC) IgG^+^) B cells and transduced the populations as described above (Figure 1B). Similar to B cell populations isolated from PBMC of healthy subjects, we again observed transduction efficiencies between 45-80% for all B cell subsets based on GFP expression (Figure 1D, F).

**Figure 1.**
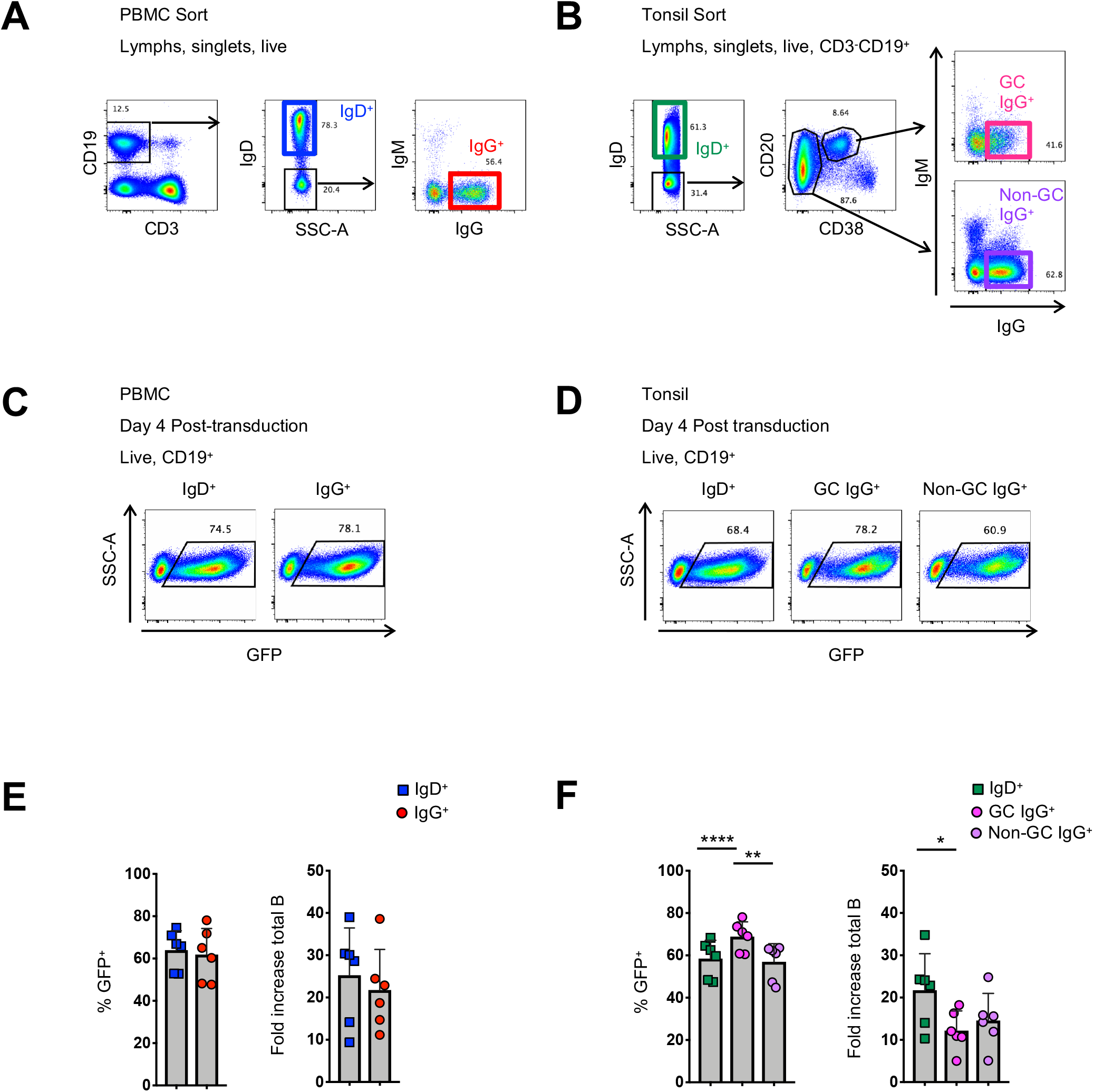
Activation and transduction of B cell subsets from PBMC and tonsil. (A, B) Flow cytometry sorting schematic for the isolation of indicated B cell populations from PBMC (A) and tonsils (B). (C, D) Representative flow plots used to determine the frequency of GFP+ cells 4 days after transduction in populations from PBMC (C) and tonsil (D). (E, F) Frequency of GFP+ B cells 4 days post-transduction (left panel) and the fold-increase in total B cell number (right panel) during activation and transduction of isolated PBMC populations (E; n=6) and of isolated tonsil populations (F; n=6). Significant differences were determined by paired t-tests for PBMC populations and one-way ANOVA with Tukey’s multiple comparisons test for tonsil populations. **** p<0.0001; **p<0.005; *p<0.05

In addition to determining the frequency of GFP^+^ B cells, we calculated the expansion of B cell subsets from the initial activation through 4 days post-transduction (6 days total). Overall, while we did observe differences between subjects, we found that the total number of CD19^+^ B cells expanded between 4-fold to 40-fold, with IgG^+^ B cells from the tonsil expanding less compared to naïve B cells (Figure 1E,F). Thus, while there are differences between B cell subsets regarding their ability to proliferate in response to CD40L and IL-21 *ex vivo*, the overall transduction efficiency of 45-80% is measured within a population of B cells that have expanded. Following transduction and expression of Bcl-6 and Bcl-xL, all GFP^+^ B cells continue to expand and to express surface immunoglobulin (Supplemental Figure 2A-C).

### Microculture of transformed B cells

To evaluate the efficiency of transformed naïve and IgG^+^ memory B cell subsets to expand in microculture, and to determine the amount of immunoglobulin secreted, we compared microcultures of transformed B cells isolated from naïve and memory B cells from both PBMC and tonsil tissue. Transduced B cells were sorted into 384-well microculture plates at 1, 2 or 5 GFP^+^ cells per well at four days post-transduction. Cultures were supplemented with fresh media, IL-21 and irradiated CD40L-feeder cells every four days. On day 24 of culture, we measured IgM and IgG concentrations in the culture supernatant of transformed naïve and memory B cells, respectively. When sorted at 1 cell per well, between 20 and 33% of the memory wells had detectable levels of IgG, while between 32 and 38% of the naive wells had detectable levels IgM (Figure 2A). Within the positive wells, the median immunoglobulin concentration ranged from 0.39 μg/mL to 1.5 μg/mL for IgG with greater median concentrations for IgM at 7.3 to 8.3 μg/mL for PBMC and tonsil cells, respectively. Sorting 2 and 5 GFP^+^ B cells per well resulted in an increased frequency of positive wells at day 24 with a slight, but not significant increase in the median immunoglobulin concentration (Figure 2A).

**Figure 2.**
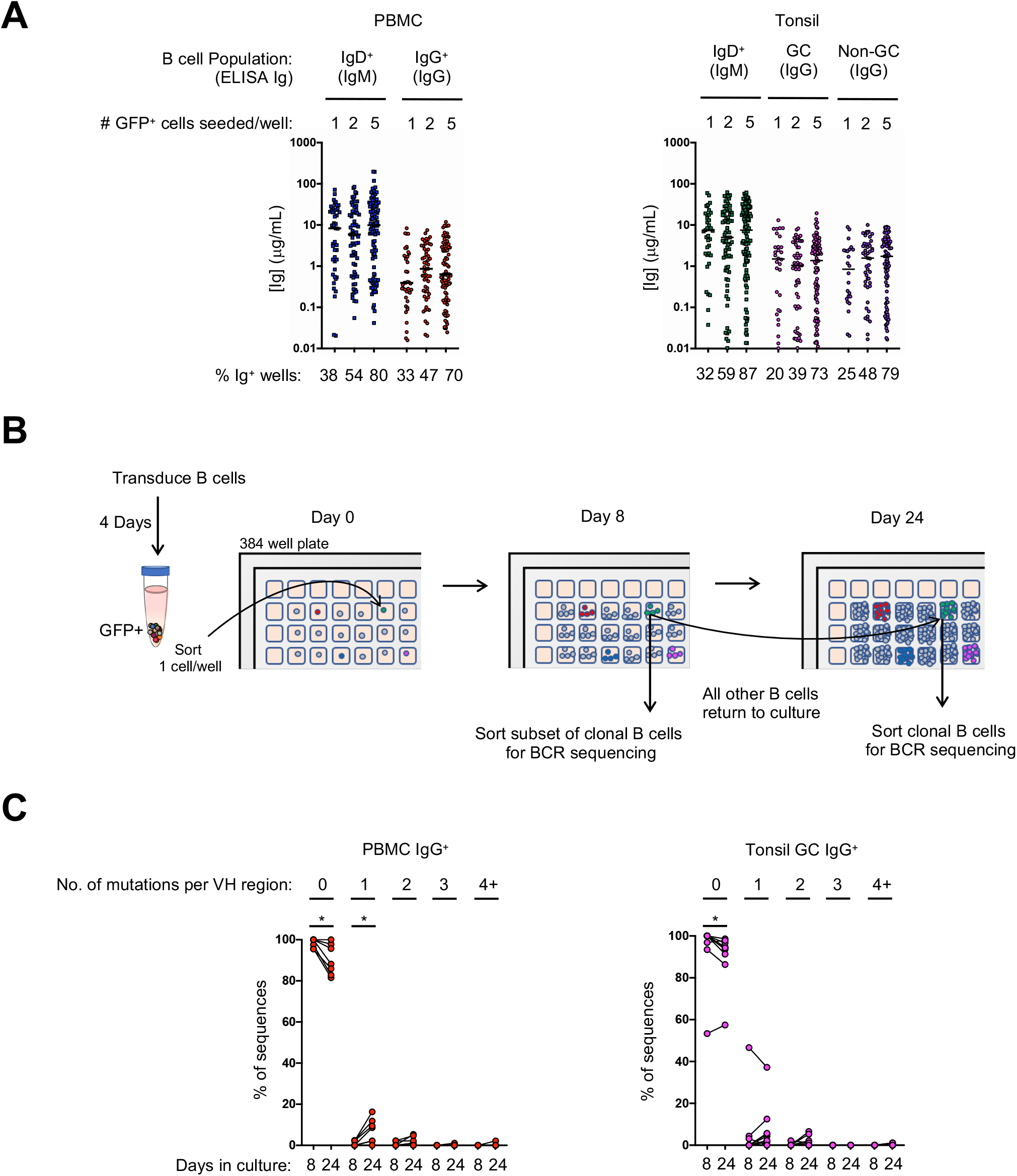
Transduced B cell populations expand and secrete immunoglobulin during 24 days of microculture. (A) Ig supernatant concentrations of IgM and IgG from PBMC (left) and tonsil (right). Horizontal bars indicate the median Ig concentration. The mean frequency of Ig positive wells is indicated at the bottom of the plots (n=3 for PBMC; n=3 for tonsil). (B) Outline of the procedure to determine the number of mutations acquired during *in vitro* culture. (C) Frequency of sequences within each clone that contained either zero, 1, 2, 3 or 4 or more mutations within the VH domain on days 8 and 24 (B: IgG+ PBMC clones, n=7; C: GC IgG+ tonsil clones, n=9). Significant differences between days 8 and 24 were determined by Wilcoxon matched-pairs signed rank test. *p<0.05

Since B cell receptor sequencing of B cells and cloning of antibodies may not occur until after screening culture supernatants on day 24 or later, we explored whether or not B cell clones acquire mutations during long-term culture, as has been described for an immortalized RSV-specific B cell line (9). We sorted single IgG^+^ transformed B cells from both PBMC and the tonsil germinal center into a microculture plate and sequenced individual B cells from the expanded clones both early during culture (day 8) and at the end of culture (day 24) (Figure 2B). Germinal center cells were specifically chosen for analysis because they express high levels of activation-induced cytidine deaminase (AID) *in vivo*, which could result in more mutations during *in vitro* culture following transformation. After aligning the clonal individual B cell sequences to determine the consensus sequence bioinformatically, we determined whether each individual sequence contained any base pair substitutions, or insertions/deletions compared to the consensus. The overwhelming majority of sequences had zero mutations at both day 8 and at day 24 of culture, although we did detect more sequences containing mutations on day 24 (PBMC IgG^+^ with zero mutations: mean 98.4% on day 8 vs 90.3% on day 24, p<0.05; Tonsil GC IgG^+^ with zero mutations: mean 93.7% on day 8 vs 90.1% on day 24, p<0.05) (Figure 2C). Interestingly, with the exception of one clone, germinal center B cells did not exhibit significantly more mutations compared to IgG^+^ B cells from PBMC. The few acquired mutations consisted mostly of one or two base pair substitutions (Figure 2C, Supplemental Figure 3A,B). These results suggest that the frequency of mutations acquired during transformed B cell culture is mostly negligible, and that the immunoglobulin sequence of the primary B cell can readily be determined by aligning multiple clonal B cell sequences to determine the consensus sequence. Thus, microculture of Bcl-6/Bcl-xL-transformed B cells is a suitable strategy to identify B cells and antibodies of interest by screening the culture supernatant for antibody function, and subsequently cloning the BCR after long-term culture.

### Generation of antigen-specific naïve and IgG^+^ memory B cell clones

We next asked whether we could generate antigen-specific memory B cell clones from both naïve and memory B cell subsets. To generate naïve B cell clones, we isolated CD27^-^IgD^+^ B cells from PBMC of a healthy donor and transduced the cells with the Bcl-6/Bcl-xL retroviral vector. Following transduction, we stained the cells with eOD-GT8, a probe that binds to VRC01-class B cells that target the CD4 binding site (27), and the eOD-GT8 knockout probe (eOD-GT8 KO), which has been mutated within the CD4 binding site and is used to avoid sorting off-target B cells (28) (Figure 3A). From the GFP^+^eOD-GT8 KO^-^ population we sorted the eOD-GT8^+^ B cells into a microculture plate at 1 cell per well. After allowing the B cells to expand, we sequenced the clonal B cells to determine their BCR sequence. Based on the sequence data we determined that clone IMMO-A1, which is of the VH1-2 family and contains a 5 amino acid long L-CDR3, is a VRC01 precursor B cell line (Figure 3B). Clone C1, which contains a VH1-2 sequence without a 5 amino acid long L-CDR3, and clone A2, which is not a member of the VH1-2 family, are also shown. After expansion and freeze/thaw, clone IMMO-A1 maintains its ability to bind to eOD-GT8.

**Figure 3.**
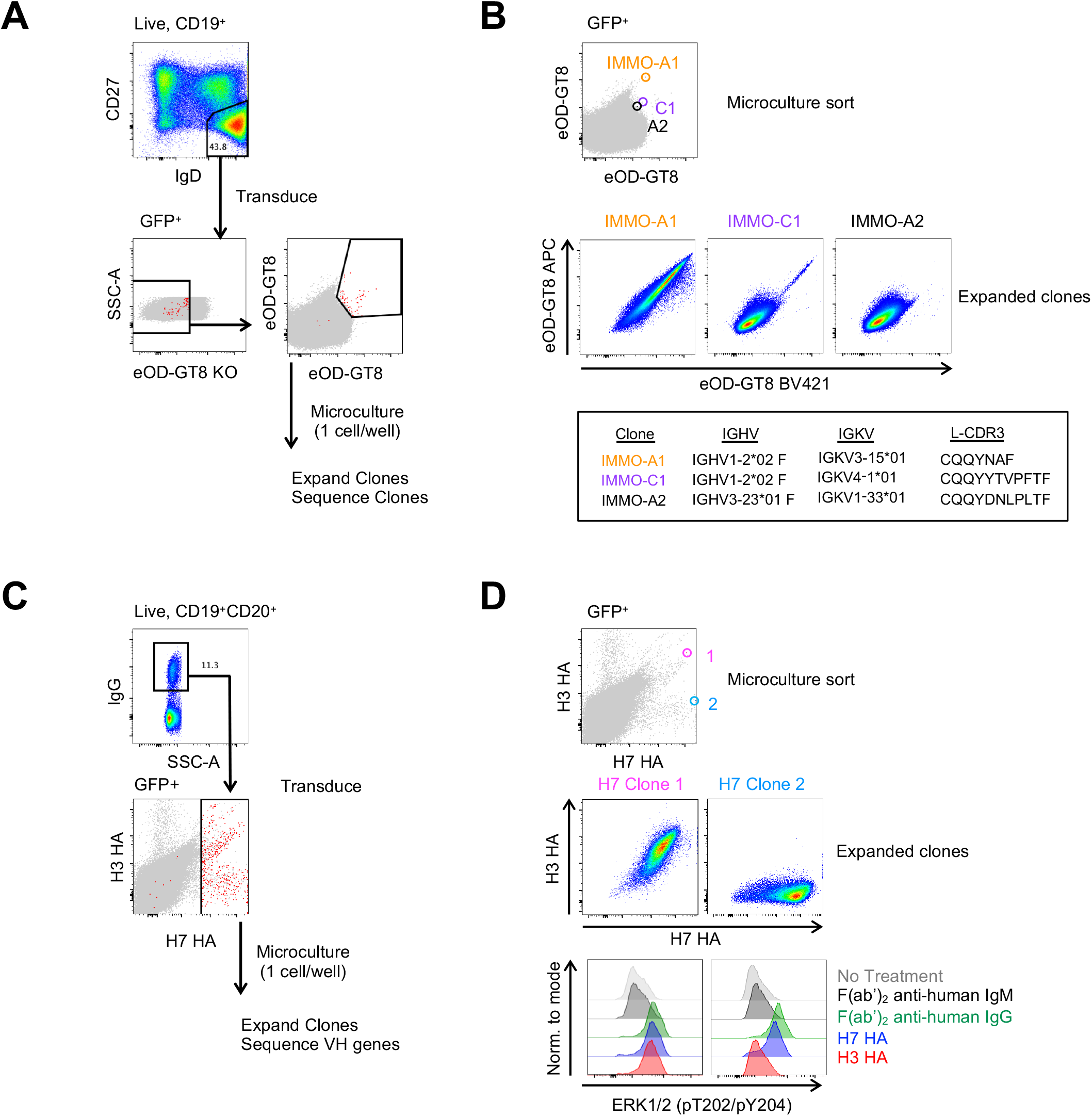
Generation of antigen-specific clonal B cell lines from naïve and memory B cell subsets. (A) Isolation of VRC01-class precursor B cell clones from the naïve B cell subset. (B) Characterization of 3 of the isolated clones binding to the eOD-GT8 probe during the microculture sort *(top)*, following clonal expansion *(middle)* and their immunogenetic properties *(bottom)*. (C) Isolation of H7 HA-specific B cells from vaccinated individuals. (D) Characterization of 2 of the isolated clones and their binding properties to the H7 and H3 HA probes (*Top:* Individual GFP^+^ B cells binding to the H7 and H3 HA probes during the microculture sort; *Middle:* binding properties to H7 and H3 HA after clonal expansion; *Lower:* BCR activation as determined by phosphorylation of ERK by flow cytometry

We also applied B cell immortalization to isolate memory B cell clones in the context of influenza vaccination, where H7N9-naïve individuals were immunized with H7N9 and subjects demonstrated B cell responses to not only the the H7 hemagglutinin (HA), but also had cross-reactive memory B cell responses to other HA molecules, such as the H3 HA. (29-31). In this context, B cell clones can be used to help characterize not only HA binding but also to demonstrate BCR activation in response to different HA molecules. To generate H7 HA-specific B cell clones we sorted IgG^+^ B cells from PBMC and following transduction we stained the transduced cells with H7 HA and H3 HA fluorophore-conjugated probes, and subsequently sorted all H7 HA-positive B cells at 1 cell per well for microculture (Figure 3C). After clonal expansion we sequenced and characterized clones for binding to H7 and H3 HA, of which Clone 1 and Clone 2 are shown (Figure 3D). We found the HA clones maintained their specificity from the single cell sort through clonal expansion, with Clone 1 binding to both H7 and H3 HA, and Clone 2 binding to the H7 HA but not to the H3 HA (Figure 3D). In addition, we found the clones maintained their specificity with respect to B cell activation as detected by the phosphorylation ERK as measured by flow cytometry, similar to what has been described for tetanus-specific B cells (9). The anti-IgG F(ab’)2 and H7 HA promoted phosphorylation of ERK in both clones, whereas the H3 HA induced phosphorylation of ERK in only Clone 1 (Figure 3D). Thus, immortalized clones can be used to demonstrate that immunogens not only bind to a specific BCR but also to activate signalling through the BCR.

Because the H7 HA-specific B cell repertoire has previously been characterized in these individuals, we evaluated how well the H7 HA-specific immortalized B cell repertoire compared to the previously characterized repertoire (30). We compared the VH gene usage of the H7 HA-specific B cells from immortalized and *ex vivo* sorted B cells from 3 donors at the same time point following vaccination (Supplemental Figure 4). In general, large clonal expansions are detected by both methods with less overlap observed with infrequent clones. Overall, while there are some differences in the VH gene usage amongst the H7 HA-specific clones, there is not a major skewing of the repertoire when comparing the *ex vivo* probe-sorted repertoire to that of the immortalized probe-sorted repertoire.

### Immortalization of B cells from individuals with chronic infection

In chronic infection settings, the CD27^-^CD21^-^ B cell population, which has a markedly reduced ability to proliferate *in vitro*, increases in frequency within PBMC. Since antigen-specific antibodies can be found within this population, we characterized the efficiency of immortalization within this population. We sorted the CD27^-^CD21^-^ and CD27^+^CD21^+^ populations from 3 healthy individuals, 3 HIV-1-viremic individuals (viral loads: 385, 4503 and 608,544 copies/mL) and 4 individuals recovering from febrile malaria at one week after treatment (convalescence). Within the HIV-1-infected and malaria convalescent donors, we observed that the *ex vivo* CD27^+^CD21^+^ population was decreased compared to healthy donors and that the CD27^-^CD21^-^ population was increased compared to healthy subjects (Figure 4A, 4B). We observed significant differences in the transduction efficiencies between the CD27^+^CD21^+^ and CD27^-^CD21^-^ population across all donors (p<0.0001) and differences across multiple subsets based on CD27 and CD21 expression (Figure 4C, Supplemental Figure 5A). We observed a similar trend with respect to the fold increase in total B cells during the transduction process, where the fold increase in B cell numbers of CD27^-^CD21^-^ population was consistently lower compared to the CD27^+^CD21^+^ population (p<0.005) (Figure 4D, Supplemental Figure 5B). On day 7 we analyzed the surface phenotype of the transduced (GFP^+^) and non-transduced (GFP^-^) populations and found that the GFP^+^ cells continued to express CD20 and surface IgG, with GFP^-^ cells losing expression of these markers (Supplemental Figure 5C). Interestingly, we found that the GFP^+^ cells expressed CD21 but little to no CD27 irrespective of the starting population. To evaluate the efficiency of B cell expansion and secretion in microculture, we sorted 2 GFP^+^ B cells per well into a 384-well plate and cultured the cells for 24 days. Although not statistically significant, we found the CD27^-^CD21^-^ population tended to result in fewer IgG^+^ positive wells compared to the CD27^+^CD21^+^ population, but that IgG^+^ wells contained similar median concentrations of IgG (Figure 4E,F, Supplemental Figure 5 D-F). Thus, while the overall B cell expansion and transduction efficiency is lower for the CD27^-^CD21^-^ population, the transduced cells do maintain surface IgG expression and secrete antibodies.

**Figure 4.**
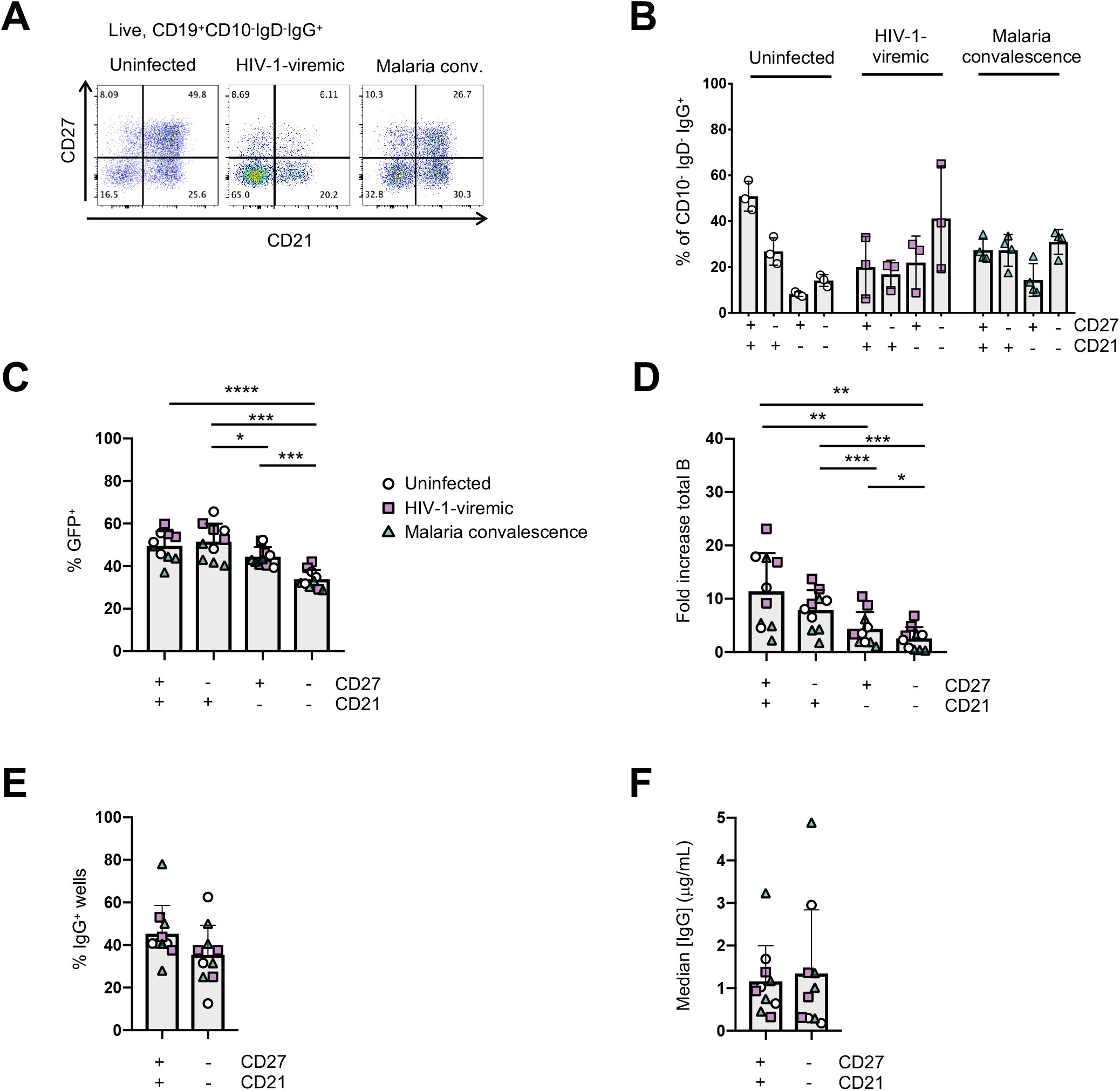
Immortalization of B cell subsets based on CD27 and CD21 expression in uninfected and HIV-1-viremic individuals and individuals treated for malaria. (A) Representative flow plots indicating the frequencies of CD27^+^CD21^+^, CD27^-^CD21^+^, CD27^+^CD21^-^ and CD27^-^CD21^-^ B cells within the IgG^+^ B cell subset from PBMC for uninfected (n=3), HIV-1-viremic (n=3) and following malaria infection (n=4). (B) Quantitation of subsets based of the expression of CD27 and CD21 within the IgG^+^ B cell subset. (C) Frequency of GFP^+^ B cells 4-days post-transduction for each isolated subset. (D) The fold-increase in total B cell number during activation and transduction for the isolated B cell populations over 6 days. (E) The frequency of wells with supernatants containing IgG was determined by ELISA after 24 days of culture. (F) The median IgG concentration within the IgG^+^ wells. Comparison between groups was performed by one-way ANOVA with significance determined by Tukey’s multiple comparisons test. ****p<0.0001; ***p<0.0005; **p<0.005; *p<0.05

In the HIV-1-infected individuals, we identified IgG-containing culture supernatants that bound to a soluble, HIV-1 envelope protein based on the BG505 sequence (BG505 SOSIP) in both the CD27^+^CD21^+^ and CD27^-^CD21^-^ populations (32) (Supplemental Figure 6A). After expanding B cells from the wells that showed binding to the BG505 SOSIP, and screening supernatants for neutralization against the BG505.T332N or the Tier1A MW965.26 pseudoviruses, we purified antibodies from 2 wells to further test for neutralizing activity (Supplemental Figure 6B). While well 1 failed to neutralize either the BG505.T332N or MW965.26 pseudoviruses upon further inspection, we did detect weak neutralization against the MW965.26 pseudovirus for well 2, which originated from the CD27^-^CD21^-^ B cell subset (Supplemental Figure 6C, D). These data demonstrate that HIV-1-specific antibodies can be isolated from the CD27^-^CD21^-^ population by B cell immortalizaton.

### Isolation of broadly neutralizing antibodies from immortalized B cells

To determine if B cell immortalization can be used to isolate bNAbs from HIV-1-infected individuals, we applied the technology to donors from which bNAbs have previously been isolated. Antibody VRC38.01 was previously isolated from donor N90 while VRC13 was isolated from patient 44 (Pt.44) (33, 34). B cells were sorted and transduced from each donor, and on day 4-post transduction GFP^+^ cells that bound to the RSC3 probe (Pt.44) and JRFL.DS.SOSIP (N90) were sorted into 384-well plates at 2 cells per well and cultured for 23 to 31 days (Figure 5A). Single point neutralization screening was performed using supernatants from days 23, 27 and 31 of microculture against the JRFL.JB pseudovirus (Figure 5B). Cell lysates from wells with greater relative neutralization were sequenced and analysis of the antibody sequences indicates the antibody isolated from donor N90 (well 5J18) is a relative of VRC38, while the antibody isolated from Pt.44 (well 3E06) is related to VRC13 (Figure 5C). Of note, the immortalization procedure was repeated for Pt.44 and a second VRC13 relative was isolated (well E3). The antibody isolated from Donor N90 (well 5J18) utilizes the same V and J genes as VRC38.01 for both the heavy and light chains and share the same CDRH3 and CDRL3 sequences. Similarly, the two antibodies isolated from Pt.44 share the same V and J genes for both the heavy and light chains as VRC13, and within the CDR3 regions residues that differ from VRC13 are highlighted in red (Figure 5C).

**Figure 5.**
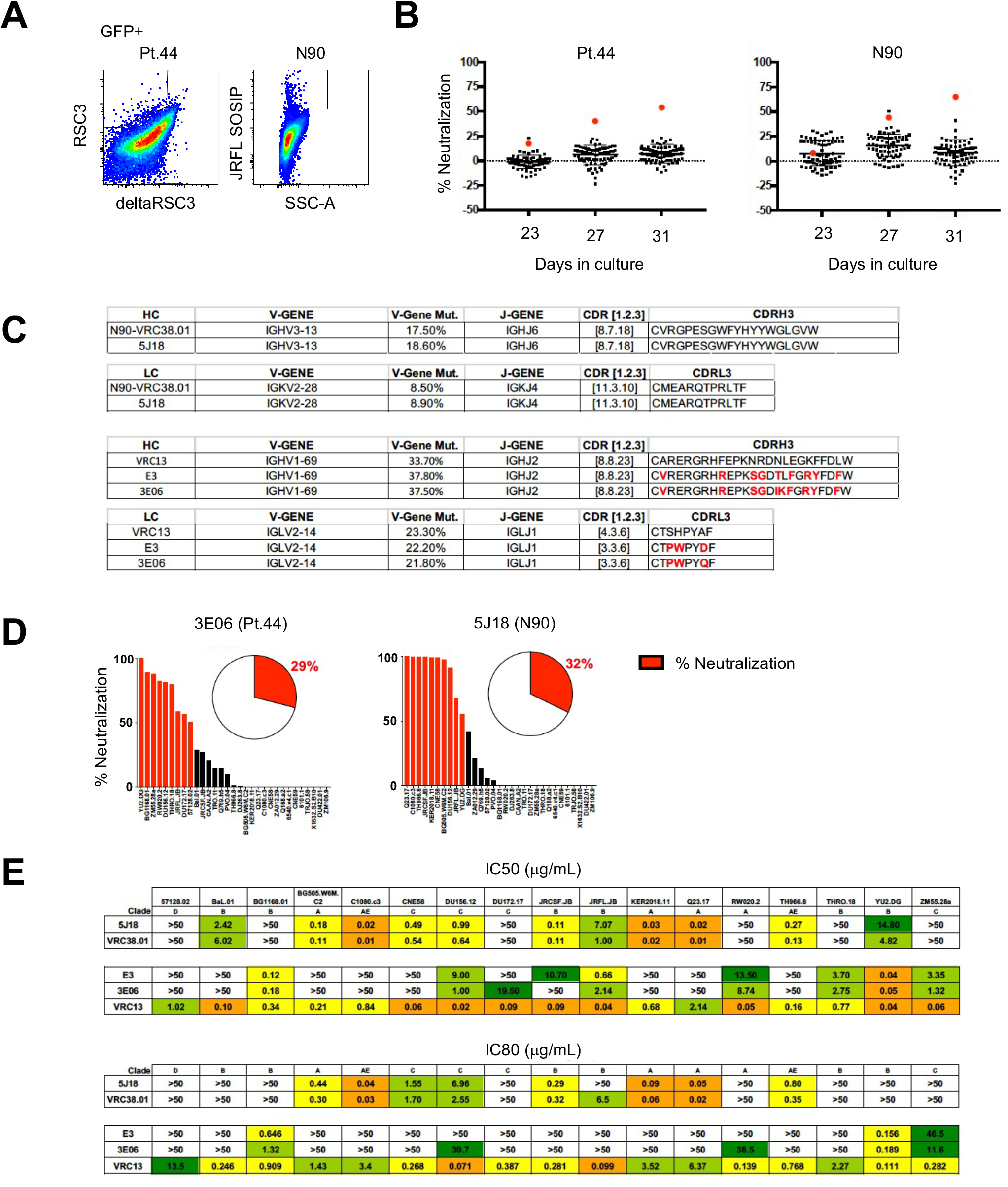
Isolation of bNAbs from HIV-1-infected individuals. (A) Flow cytometry plots showing B cell binding to HIV-1-envelope specific probes after B cell transduction. (B) Percent neutralization of B cell culture supernatants against the JRFL pseudovirus following 23, 27 or 31 days of microculture. (C) Immunogenetic characterization of antibodies sequenced from wells identified as having greater relative neutralization. (D) Single point neutralization assays were performed against a 31-virus panel for the cloned antibodies. (E) IC50 and IC80 values were determined for the three identified antibodies using a panel of 17 pseudoviruses.

Single point neutralization assays were performed against a 31-virus panel for the three cloned antibodies and between 29 and 32% of viruses were neutralized by more that 50% (Figure 5D). The cloned antibodies, along with VRC13 and VRC38.01 were further tested for neutralization against a panel of 17 viruses that showed sensitivity in the single-point assay. The antibody isolated from well 5J18 showed similar potency and breadth as compared to VRC38.01, whereas the two antibodies isolated from Pt.44 display less potency and breadth as compared to VRC13 (Figure 5E). Overall, these studies serve as a proof-of-principle that B cell immortalization can be used to isolate bNAbs.

## Discussion

While Bcl6/Bcl-xL B cell immortalization was first described in IgG^+^ and IgM^+^ memory B cell populations in PBMC from healthy donors, and similar B cell immortalization methods have recently been described in cancer settings and applied to naïve B cells, an extensive comparison of the same Bcl6/Bcl-xL B cell immortalization protocol across multiple B cell subsets, including chronic infection settings, has not been done (9, 27, 35). Here, we characterized and applied Bcl-6/Bcl-xL B cell immortalization to multiple B cell subsets including naïve and memory B cell subsets from both lymphoid tissue and PBMC, and in the CD27^-^CD21^-^ and CD27^+^CD21^+^ populations in chronic infection. In our hands, we found Bcl-6/BclxL B cell immortalization to be applicable across multiple B cell subsets, although with differing efficiencies amongst distinct B cell populations.

The application of B cell immortalization to B cell populations with a reduced capacity to proliferate and survive *in vitro*, such as CD27^-^CD21^-^ memory B cells and germinal center B cells, may provide an alternative means for antibody isolation from these populations, in lieu of traditional B cell culture. For example, although we found primary germinal B cells expanded less when cultured with IL-21 and CD40 Ligand when compared to other B cell subsets, we found that post-transduction, the immortalized B cells from this population expanded and secreted antibodies in similar amounts to that of immortalized IgG^+^ B cells from PBMC of healthy donors, suggesting that immortalization of germinal center B cells could serve as an alternative method to isolate rare antibodies from germinal centers of SLOs. In this regard, HIV-1-specific bNAbs were recently isolated from germinal center B cells of lymph nodes using an Epstein-Barr Virus-based immortalization strategy, demonstrating that rare antibodies can be isolated from germinal centers with B cell immortalization strategies (36). In chronic infection settings, we found that B cell immortalization can be applied to the CD27^+^CD21^+^ population with similar efficiency to that seen in IgG^+^ memory B cells in uninfected individuals, with decreased transduction and expansion in the CD27^-^CD21^-^ population. Thus, if antibodies of interest are found within the CD27^+^CD21^+^ population, this technique has a greater capacity to isolate the antibodies of interest, although antibodies can still be isolated from the CD27^-^ CD21^-^ population, as the immortalized B cells from this population do secrete antibodies and express surface immunoglobulin.

One of the greatest advantages of this technology is the flexibility of screening for both antibody binding and function with relatively large quantities of immunoglobulin-containing culture supernatant. In a traditional B cell culture method, where primary B cells are activated for 10 to 14 days before screening immunoglobulin-containing culture supernatant, a limited amount of functional screening can be applied. In the HIV-1 TZM-bl microneutralization assay, which screens primary B cell culture supernatants from 384-well culture plates for HIV-1 neutralizing antibodies, one may only test for neutralizing activity against one or two pseudoviruses due to limited supernatant volumes. Because immortalized B cells are continuously expanding, larger volumes of supernatant can be generated and screened against several pseudoviruses to assay for breadth before antibody sequencing and cloning. We were able to isolate a HIV-1-specific antibody from the CD27^-^CD21^-^ population of a HIV-1-infected donor, where we first screened for binding to the HIV-1 envelope SOSIP trimer, with subsequent screening for neutralizing activity after expanding wells of interest. Importantly, we were also able to demonstrate this technology can be applied to isolate bNAbs from HIV-1-infected indiviudals by using a slightly different strategy where we combined B cell immortalization with the TZM-bl microneutralization assay and subsequently cloned the antibodies for further characterization. Both of these examples, along with our recent study in donors after malaria infection where we isolated MSP-specific IgM^+^ antibodies, can serve as a proof-of-concept to show that Bcl6/Bcl-xL immortalization can be applied to chronic infection settings to isolate antibodies of interest (37).

Because we did detect differences between B cell subsets during activation and following transduction, and because individual clones do expand at different rates, the utility of this methodology as a means to characterize the B cell repertoire is unclear. Using influenza vaccination as a model system, we compared the H7 HA-immortalized B cell repertoire to that of a previously characterized repertoire using the same samples and the same probes as bait as previously published (30). In general, both methodologies detected large clonal expansions but there was less overlap with rarer clones, with some clones detected with only *ex vivo* sorting, and others detected after immortalization and clonal expansion. And while some of these discrepancies can be attributed to the depth of sequencing, they may also be attributed to the variable efficiency of B cell immortalization in different B cell subsets. In general however, we did not detect a major bias in the immortalized H7 HA-specific B cell repertoire compared to the previously published *ex vivo* repertoire (30). Therefore, while B cell immortalization can be used to isolate rare antibodies and generate B cell clones, its application to accurately study the B cell repertoire needs to be further explored.

As B cell clones are expanded for increased screening and analysis, one potential concern is whether the clones acquire mutations during the *in vitro* culture, and whether the final sequence of the desired clone represents the sequence of the primary B cell. In our experimental setting, while we did detect mutations during the microculture, we found these mutations to be minimal. Some of the mutations may be due to PCR error, although some certainly represent actual mutations as we observed more mutations on day 24 of culture compared to day 8 using the same sequencing protocol. Overall, we found that by sequencing several individual B cells from the same clone and aligning the sequences bioinformatically, we could determine the consensus sequence which likely represents the sequence of the original primary B cell. Of course, as clones are further expanded one should take into account the accumulation of additional mutations which may or may not be desired.

An additional advantage of using immortalization technology is the generation of clonal B cell lines, which can serve as valuable tools in the development of new reagents and assays. As novel antigens are developed to serve as probes for flow cytometric isolation and characterization of antigen-specific B cells, immortalized B cell clones can serve as tools to characterize the binding specificity and potential competition between related probes. Could cross-reactive B cells be missed if related probes used within the same flow cytometry panel compete for binding to the same B cell clonal family? To better understand this, B cell clones can be used in to screen related probes for competitive binding to BCRs of interest.

In addition, immortalized B cell clones can serve as tools to characterize the capacity of novel immunogens to not only bind to antigen-specific B cells but also to induce B cell signaling. In combination with a high-throughput B cell activation assay, such as detection of phosphorylation of ERK by flow cytometry, libraries of immortalized B cell clones can be utilized to measure the diversity and specificity of the B cells that respond to the immunogen. The use of clonal B cell lines may also prove to be especially valuable in the HIV-1 vaccine field where novel immunogens are being designed to bind to and activate germline precursor B cells. Here, naïve precursor clonal B cell lines can serve as controls to characterize novel probes aimed to activate precursor B cells and to also test for assay specificity, sensitivity and reproducibility in the analysis of data from clinical trials.

As with any methodology, there are advantages and disadvantages to be considered before applying the technology, including the efficiency of the methodology. Overall, Bcl-6/Bcl-xL B cell immortalization is a valuable tool for antibody isolation and clonal B cell line development, and our results here show that it can be applied to different B cell populations, including populations that may be less amenable to long-term B cell culture such as in chronic infection settings.

## Materials and Methods

### Human Subjects

Peripheral blood mononuclear cells (PBMC) from healthy subjects were obtained from donors participating in the NIH research apheresis program. Tonsil cells were acquired from discarded anonymized specimens from Children’s National Medical Center (CNMC) with the approval from the Basic Science Core of the District of Columbia Developmental Center for AIDS Research and did not constitute ‘human subjects research’ as determined by the CNMC Institutional Review Board. Samples from the H7N9 influenza vaccine study (VRC 315; ClinicalTrials.gov; NCT02206464), which was a randomized phase I clinical trial in healthy adults designed to study the safety and immunogenicity of prime-boost vaccination regimens, were obtained through the VRC clinic (29-31). Informed consent was obtained from each volunteer and approved by the Institutional Review Board at NIAID, NIH. PBMC samples from three HIV-1-infected donors were also obtained through the Vaccine Research Clinic, while PBMC from HIV-1-infected individuals with broadly neutralizing sera (patient 44 and donor N90) have been described previously (33, 34). All donors provided informed consent and studies were approved by the Institutional Review Board at NIAID, NIH. PBMC samples collected one week after treatment of the first febrile malaria episode were obtained from a Malian cohort as previously described (38), and was approved by the Ethics Committee of the Faculty of Medicine, Pharmacy and Dentistry at the University of Sciences, Technique, and Technology of Bamako, and the Institutional Review Board of NIAID, NIH (ClinicalTrials.gov; NCT0132258).

### Reagents

pLZRS-IRES-GFP was kindly provided by Lynda Chin (Addgene plasmid # 21961; http://n2t.net/addgene:21961) (39). The human codon-optimized Bcl6-P2A-human codon-optimized Bcl-xL insert was synthesized using GeneArt (ThermoFisher) and cloned into pLZRS-IRES-GFP. 3T3-msCD40L cells were cultured and irradiated (5000 rads) as described previously (8). GP2-293 retroviral packaging cells and the p10A1 envelope vector were purchased from Clontech. Lipofectamine 3000 was purchased from ThermoFisher and Polybrene was purchased from Sigma. Cell culture reagents, including DMEM, IMDM, 100X Penicillin/Streptomycin/Glutamine were purchased from Gibco. Probes to detect antigen specific B cells against influenza (H7 and H3 HA), CD4-binding site antibodies against HIV-1 (RSC3 and deltaRSC3) and to detect VRC01-class precursor B cells (eOD-GT8 and eOD-GT8 knockout monomers) were generated and conjugated to fluorochromes as previously described (27, 28, 30, 40, 41).

### Human IL-21 production

Human IL-21 was produced at the Vaccine Research Center. To express human interleukin-21 (IL21), (UniProtKB Q9HBE4) with a C-terminal thrombin cleavage site, an 8xHisTag and a TwinStrep tag the gene was synthesized and cloned into a mammalian expression vector pVRC8400. The plasmid was transiently transfected into freeStyle293F cells (Thermo Fisher). Protein was purified from filtered cell culture supernatant by Ni-NTA affinity column first and then by a StrepTactin (IBA) column.

### B cell staining and sorting

To sort B cell populations from PBMC and tonsil cells (Figure 1), cells were first stained with violet fluorescent dye (Invitrogen) to demarcate dead cells, and then surface stained with an antibody cocktail. The antibodies used include: CD3 H7APC (clone SK7; BD Biosciences), IgM PE-CF594 (clone G20-127; BD Biosciences), IgG PE-Cy5 (clone G18-145; BD Biosciences), CD20 BV570 (clone 2H7; Biolegend), CD19 BV785 (clone HIB19; Biolegend), IgD PE (Southern Biotech) and CD38 Alexa680 (clone OKT10; conjugated at the VRC). Sorted bulk B cell populations were activated and transduced. To isolate VRC01 precursor B cells, naïve cells (CD27^-^IgD^+^) were first stained with violet fluorescent dye and subsequently incubated with antibodies CD19 BV785, CD27 PC5 (clone 1A4CD27; Beckman Coulter), IgD PE and IgG BV605 (clone G18-145; BD Biosciences). Four days following transduction cells were stained with eOD-GT8 probes (labeled with APC or BV421) and the knockout eOD-GT8 probe (BV605) and GFP^+^eOD-GT8^+^ cells were sorted into microplates containing irradiated CD40L feeder cells and IL-21 at 1 cell per well for clonal expansion and sequencing. To generate influenza-specific B cell clones, PBMC were stained with violet fluorescent dye and CD3 BV785, CD19 ECD (clone J3-119; Beckman Coulter), CD20 APC-Cy7 (clone 2H7; Biolegend) and IgG Alexa700 (G18-145; BD Biosciences), sorted, transduced and subsequently stained with CD19 ECD and fluorochrome-conjugated probes to detect H3 HA and/or H7 HA specific B cells. GFP^+^HA^+^ B cells were sorted into microculture plates at 1 cell per well and cultured as described above. To compare B cell immortalization efficiency in B cell subsets from HIV-1-infected donors and donors recovering from malaria, cells were first surface stained, sorted in bulk, transduced and GFP^+^ B cells were sorted into microculture plates at 2 cells per well. Antibodies used for surface staining include CD20 APC-Cy7, CD19 BV785, IgD BV605 (clone IA6, BD Biosciences), CD10 PE-Cy5 (clone HI10a; BD Biosciences), IgG Alexa700, CD27 BV650 (clone 0323; Biolegend) and CD21 PE-Cy7 (clone B-ly4; BD Biosciences). All surface stains were incubated with the appropriate antibody cocktails for 20 minutes at room temperature and B cells were sorted using a modified FACSAria. Flow cytometry data were analyzed in Flowjo v 10.8.1 or v 9.9.6 (TreeStar).

### B cell isolation and activation

B cell populations from PBMC or tonsils were sorted as described above and cultured in 12-well tissue culture plates in 1 mL of I10 medium (IMDM (Gibco) with glutamine, 10% FBS, supplemented with Penicillin, Streptomycin, Glutamine). Wells contained 1 × 10^4^ to 2 × 10^5^ B cells, 2 × 10^5^ irradiated 3T3-msCD40L cells and 25 ng/mL human IL-21. Cells were cultured for 36 to 48 hours at 37°C and 5% CO2 before retroviral transduction.

### Pseudotyped retrovirus generation

5 × 10^6^ GP2-293 packaging cells (Clontech) were plated in a 10 cm dish containing D10 medium (DMEM supplemented with 10% FBS, penicillin, streptomycin and glutamine). 24 hours after plating, 25 μg of the retroviral expression vector (pLZRS-human codon-optimized Bcl6-P2A-human codon-optimized BclxL-IRES-GFP) and 5 μg of the envelope vector (p10A1; Clontech) were transfected into GP2-293 cells using Lipofectamine 3000 (ThermoFisher) according to the manufacturer’s instructions. After 24 hours, the cells were supplemented with fresh D10 media. Retroviral supernatant was harvested 48 hours post-transfection, replaced with fresh D10 and again harvested 72 hours post-transfection.

### Retroviral Transduction of Activated B cells

B cells that had been activated for 36-48 hours were transduced with fresh viral supernatant containing polybrene (4 μg/mL). Briefly, 1 mL I10 media was removed from the well of the 12-well plate and replaced with 1 mL fresh viral supernatant (48-hour retroviral supernatant) containing polybrene. The plate was centrifuged at 1200*xg* for 1 hour at 32°C, rested for 1 hour at 37°C and 5% CO2 before the viral supernatant was replaced with fresh I10 containing 25 ng/mL IL-21 and cultured at 37°C and 5% CO2 overnight. The same procedure was repeated the following day using the 72-hour retroviral supernatant. B cells were then rested for ∼3 days before determining transduction efficiency by flow cytometry and sorting for microculture.

### Transformed B cell culture

Transformed B cells cultured in bulk were maintained in 12-well plates. Cells were split 1:4 every 3-4 days into wells containing 2 ×10^5^ 3T3-msCD40L cells and 25 ng/mL human IL-21 in 1mL of I10. Cell counts were measured by flow cytometry using counting beads (BD Biosciences) and data were collected on a modified LSRII or a modified FACSymphony. For microculture, transformed B cells were sorted 3 days following transduction. Briefly, transformed B cells were sorted at 1-5 B cells/well into a 384-well plate containing 5 × 10^3^ irradiated 3T3-msCD40L, human IL-21 (25 ng/mL) in 50 μL of I10. Every 4 days cells were fed by replacing 25 μl of media with fresh I10 containing irradiated 2.5 × 10^3^ 3T3-msCD40L cells and 50 ng/mL IL-21 using a Biomek NX^P^ liquid handler (Beckman Coulter). Cells were cultured for 2-3 weeks before harvesting the final supernatant and either freezing the cultures in the 384-well plate (50ul freezing media per well (FBS + 10% DMSO)), or expanding selected wells to 96-well tissue culture plates.

### IgG and IgM ELISA

Concentrations of total IgG and IgM in the culture supernatant were determined by ELISA (ThermoFisher) according to the manufacturer’s instruction.

### Immunoglobulin amplification and sequencing

B cells were dry-sorted into a 96-well PCR plate at 1-2 cells per well and multiplex polymerase chain reaction (PCR) was used to amplify the heavy and/or light chain sequences as previously described (7). PCR products were sequenced by ACGT, Inc., Eurofins or Genewiz and analyzed using Geneious Prime and the IMGT database (42, 43).

### In vitro mutation analysis

To characterize the accumulation of mutations during *in vitro* culture, IgG^+^ B cells from PBMC of uninfected individuals and germinal center IgG^+^ B cells from tonsil tissue were transduced. Four days post-transduction, GFP^+^ B cells were sorted at one cell per well into 384-well microculture plates. Following 8 days of microculture, clonal GFP^+^ B cells from selected wells were harvested and sorted at 1 cell per well into a PCR plate for BCR sequencing (45 total wells for each clone), and the remaining clonal B cells were placed back into culture. The same selected clonal B cells were again harvested on day 24 and sorted into PCR plates at 1 cell per well for BCR sequencing (93 total wells for each clone). The sequences from the VH genes were aligned using Geneious software to generate a consensus sequence and each individual sequence was analyzed for differences from the consensus sequence. Mutations deviating from the consensus sequence were confirmed by analyzing sequence chromatograms and assessing the quality of chromatogram peaks relative to the basecall recorded. Only mutations with clear, single color, high confidence sequence peaks were considered as actual mutations.

### Phosphoflow to measure phosphorylation of ERK

HA-specific immortalized B cell clones were resuspended in IMDM + 1% FBS and stimulated for 2 minutes at 37°C with HA probes (250nM), anti-IgG F(ab’)2 (5 μg/mL; Southern Biotech) or anti-IgM F(ab’)2 (5 μg/mL; Southern Biotech). Cells were immediately fixed with pre-warmed paraformaldehyde (4% final concentration), incubated at 37°C for 10 minutes and pelleted by centrifugation. Cells were resuspended in cold Perm Buffer III (BD Biosciences), incubated on ice for 30 minutes and washed twice with stain buffer (PBS + 1% FBS). Cells were resuspended in stain buffer and incubated with anti-ERK1/2 (pT202/pY204 Alexa647) (BD Biosciences) for 30 minutes at room temperature. Cells were subsequently washed and data were collected on a modified LSRII (BD Immunocytometry Systems) and analyzed using Flowjo (TreeStar).

### BG505-SOSIP DS Binding Analysis of Supernatants

Standard MSD 384 well streptavidin coated SECTOR®Imager 2400 plates were blocked with 35 µL of 5% (W/V) MSD Blocker A and incubated for 1 hour at room temperature (RT) on a Heidolph Titramax 100 vibrational plate shaker at 650 rpm. All incubations in this assay were performed as described above. The plates were washed three times with 0.05%Tween PBS (wash buffer) and were coated with biotinylated BG505 SOSIP DS-FPV1 10ln QQ AVI protein (Biotin Trimer 7070 capture) at an optimized concentration of 1 μg/mL for 1 hour. 1% MSD Blocker A was used as the diluent in the assay. The test samples (B cell supernatants) were diluted 1 to 1 with assay diluent in dilution plates. After 1 hour of incubation, the plates were washed again with the wash buffer and the serial diluted test samples were added to MSD plates. After an hour of incubation with the samples, the plates were washed again, and SULFO-TAG conjugated goat anti-human secondary detection antibody was used for detection at an optimized concentration of 1 μg/mL. After an additional hour of incubation, the unbound secondary detection antibody was washed off the plates and the plates were read using 1X MSD Read Buffer on the MSD Sector Imager 2400.

### Antibody purification from B cell culture supernatants

B cells from the CD27^+^CD21^+^ and CD27^-^CD21^-^ populations that displayed greater relative neutralization against pseudoviruses expressing HIV-1 envelopes BG505.T32N and MW965.26 in the TZM-bl microneutralization assay were expanded in serum free media hybridoma media (Gibco) that was supplemented with irradiated feeder cells and IL-21. Supernatants were concentrated and IgG was purified using Ab Spintrap columns (GE Healthcare/Cytiva).

### HIV-1 TZM-bl Neutralization Assays

Screening small volumes of B cell culture supernatants for HIV-1 neutralizing activity was performed using the HIV-1 microneutralization assay as described (44). B cells from Pt.44 and donor N90 that displayed greater relative neutralization against the JRFL.JB pseudovirus in the microneutralization assay were cloned and expressed. Percent neutralization was calculated as: [((RLU virus with no Ab) – (RLU virus with Ab))/(RLU virus with no Ab)] X 100. Neutralization breadth and IC50 and IC80 values were determined for HIV-1-specific antibodies against a panel of Env-pseudoviruses using TZM-bl target cells as previously described (45).

### Statistical Analysis

GraphPad Prism software was used for statistical analysis and to prepare figures. Statistical analysis was performed using paired t-tests, Wilcoxon matched-pairs signed rank tests or one-way ANOVA with Tukey’s multiple comparison test as indicated. Bars depict mean values with standard deviations shown.

## Acknowledgements

We would like to thank Drs. Nicole Doria-Rose and Rosemarie Mason for B cell culture advice and training on the liquid handling robots, and Dr. Guillame Stewart-Jones for providing the JRFL.DS.SOSIP. We would also like to thank Alida Taylor and Carmelo Chiedi for irradiating the 3T3-msCD40L cells and Sijy O’Dell and Steve Schmidt for providing the VRC01 antibody and the pSG3deltaEnv vector, and for the plasmids encoding the BG505.T332N and MW965.26 HIV-1 envelopes. We thank the VRC Flow Cytometry Facility as well as Margaret Beddall for help with flow cytometry instruments and in-house antibody conjugation. The pLZRS-IRES-GFP plasmid was kindly provided by Lynda Chin. This work was supported by the Bill and Melinda Gates Foundation (grant OPP1147555) and by the Intramural Research Program of the Vaccine Research Center, NIAID, NIH.

## Notes

### Competing Interest Statement

The authors have declared no competing interest.

## References

1. The Antibody Society: Therapeutic monoclonal antibodies approved or in review in the EU or US. [Available from: http://www.antibodysociety.org.

2. Julg B, Barouch D. Broadly neutralizing antibodies for HIV-1 prevention and therapy. Semin Immunol. 2021;51:101475.

3. Stephenson KE, Barouch DH. Broadly Neutralizing Antibodies for HIV Eradication. Curr HIV/AIDS Rep. 2016;13(1):31–7.

4. Mahomed S, Garrett N, Baxter C, Abdool Karim Q, Abdool Karim SS. Clinical Trials of Broadly Neutralizing Monoclonal Antibodies for Human Immunodeficiency Virus Prevention: A Review. J Infect Dis. 2021;223(3):370–80.

5. Wilson PC, Andrews SF. Tools to therapeutically harness the human antibody response. Nat Rev Immunol. 2012;12(10):709–19.

6. Scheid JF, Mouquet H, Feldhahn N, Walker BD, Pereyra F, Cutrell E, et al. A method for identification of HIV gp140 binding memory B cells in human blood. J Immunol Methods. 2009;343(2):65–7.

7. Tiller T, Meffre E, Yurasov S, Tsuiji M, Nussenzweig MC, Wardemann H. Efficient generation of monoclonal antibodies from single human B cells by single cell RT-PCR and expression vector cloning. J Immunol Methods. 2008;329(1-2):112–24.

8. Huang J, Doria-Rose NA, Longo NS, Laub L, Lin CL, Turk E, et al. Isolation of human monoclonal antibodies from peripheral blood B cells. Nat Protoc. 2013;8(10):1907–15.

9. Kwakkenbos MJ, Diehl SA, Yasuda E, Bakker AQ, van Geelen CM, Lukens MV, et al. Generation of stable monoclonal antibody-producing B cell receptor-positive human memory B cells by genetic programming. Nat Med. 2010;16(1):123–8.

10. Kwakkenbos MJ, Bakker AQ, van Helden PM, Wagner K, Yasuda E, Spits H, et al. Genetic manipulation of B cells for the isolation of rare therapeutic antibodies from the human repertoire. Methods. 2014;65(1):38–43.

11. Kwakkenbos MJ, van Helden PM, Beaumont T, Spits H. Stable long-term cultures of self-renewing B cells and their applications. Immunol Rev. 2016;270(1):65–77.

12. Liu YJ, Mason DY, Johnson GD, Abbot S, Gregory CD, Hardie DL, et al. Germinal center cells express bcl-2 protein after activation by signals which prevent their entry into apoptosis. Eur J Immunol. 1991;21(8):1905–10.

13. Moir S, Ho J, Malaspina A, Wang W, DiPoto AC, O’Shea MA, et al. Evidence for HIV-associated B cell exhaustion in a dysfunctional memory B cell compartment in HIV-infected viremic individuals. J Exp Med. 2008;205(8):1797–805.

14. Weiss GE, Crompton PD, Li S, Walsh LA, Moir S, Traore B, et al. Atypical memory B cells are greatly expanded in individuals living in a malaria-endemic area. J Immunol. 2009;183(3):2176–82.

15. Moir S, Fauci AS. B-cell exhaustion in HIV infection: the role of immune activation. Curr Opin HIV AIDS. 2014;9(5):472–7.

16. Portugal S, Obeng-Adjei N, Moir S, Crompton PD, Pierce SK. Atypical memory B cells in human chronic infectious diseases: An interim report. Cell Immunol. 2017;321:18–25.

17. Holla P, Ambegaonkar A, Sohn H, Pierce SK. Exhaustion may not be in the human B cell vocabulary, at least not in malaria. Immunol Rev. 2019;292(1):139–48.

18. Portugal S, Tipton CM, Sohn H, Kone Y, Wang J, Li S, et al. Malaria-associated atypical memory B cells exhibit markedly reduced B cell receptor signaling and effector function. Elife. 2015;4.

19. Kardava L, Moir S, Shah N, Wang W, Wilson R, Buckner CM, et al. Abnormal B cell memory subsets dominate HIV-specific responses in infected individuals. J Clin Invest. 2014;124(7):3252–62.

20. Muellenbeck MF, Ueberheide B, Amulic B, Epp A, Fenyo D, Busse CE, et al. Atypical and classical memory B cells produce Plasmodium falciparum neutralizing antibodies. J Exp Med. 2013;210(2):389–99.

21. Mascola JR, Haynes BF. HIV-1 neutralizing antibodies: understanding nature’s pathways. Immunol Rev. 2013;254(1):225–44.

22. Rusert P, Kouyos RD, Kadelka C, Ebner H, Schanz M, Huber M, et al. Determinants of HIV-1 broadly neutralizing antibody induction. Nat Med. 2016;22(11):1260–7.

23. Abela IA, Kadelka C, Trkola A. Correlates of broadly neutralizing antibody development. Curr Opin HIV AIDS. 2019;14(4):279–85.

24. Stamatatos L, Pancera M, McGuire AT. Germline-targeting immunogens. Immunol Rev. 2017;275(1):203–16.

25. McGuire AT. Targeting broadly neutralizing antibody precursors: a naive approach to vaccine design. Curr Opin HIV AIDS. 2019;14(4):294–301.

26. Jardine J, Julien JP, Menis S, Ota T, Kalyuzhniy O, McGuire A, et al. Rational HIV immunogen design to target specific germline B cell receptors. Science. 2013;340(6133):711–6.

27. Jardine JG, Kulp DW, Havenar-Daughton C, Sarkar A, Briney B, Sok D, et al. HIV-1 broadly neutralizing antibody precursor B cells revealed by germline-targeting immunogen. Science. 2016;351(6280):1458–63.

28. Sok D, Briney B, Jardine JG, Kulp DW, Menis S, Pauthner M, et al. Priming HIV-1 broadly neutralizing antibody precursors in human Ig loci transgenic mice. Science. 2016;353(6307):1557–60.

29. DeZure AD, Coates EE, Hu Z, Yamshchikov GV, Zephir KL, Enama ME, et al. An avian influenza H7 DNA priming vaccine is safe and immunogenic in a randomized phase I clinical trial. NPJ Vaccines. 2017;2:15.

30. Andrews SF, Joyce MG, Chambers MJ, Gillespie RA, Kanekiyo M, Leung K, et al. Preferential induction of cross-group influenza A hemagglutinin stem-specific memory B cells after H7N9 immunization in humans. Sci Immunol. 2017;2(13).

31. Andrews SF, Chambers MJ, Schramm CA, Plyler J, Raab JE, Kanekiyo M, et al. Activation Dynamics and Immunoglobulin Evolution of Pre-existing and Newly Generated Human Memory B cell Responses to Influenza Hemagglutinin. Immunity. 2019;51(2):398–410 e5.

32. Sanders RW, Derking R, Cupo A, Julien J-P, Yasmeen A, de Val N, et al. A Next-Generation Cleaved, Soluble HIV-1 Env Trimer, BG505 SOSIP.664 gp140, Expresses Multiple Epitopes for Broadly Neutralizing but Not Non-Neutralizing Antibodies. PLOS Pathogens. 2013;9(9):e1003618.

33. Cale EM, Gorman J, Radakovich NA, Crooks ET, Osawa K, Tong T, et al. Virus-like Particles Identify an HIV V1V2 Apex-Binding Neutralizing Antibody that Lacks a Protruding Loop. Immunity. 2017;46(5):777–91 e10.

34. Zhou T, Lynch RM, Chen L, Acharya P, Wu X, Doria-Rose NA, et al. Structural Repertoire of HIV-1-Neutralizing Antibodies Targeting the CD4 Supersite in 14 Donors. Cell. 2015;161(6):1280–92.

35. Caeser R, Di Re M, Krupka JA, Gao J, Lara-Chica M, Dias JML, et al. Genetic modification of primary human B cells to model high-grade lymphoma. Nat Commun. 2019;10(1):4543.

36. Pinto D, Fenwick C, Caillat C, Silacci C, Guseva S, Dehez F, et al. Structural Basis for Broad HIV-1 Neutralization by the MPER-Specific Human Broadly Neutralizing Antibody LN01. Cell Host Microbe. 2019;26(5):623–37 e8.

37. Hopp CS, Sekar P, Diouf A, Miura K, Boswell K, Skinner J, et al. Plasmodium falciparum-specific IgM B cells dominate in children, expand with malaria, and produce functional IgM. J Exp Med. 2021;218(4).

38. Tran TM, Li S, Doumbo S, Doumtabe D, Huang CY, Dia S, et al. An intensive longitudinal cohort study of Malian children and adults reveals no evidence of acquired immunity to Plasmodium falciparum infection. Clin Infect Dis. 2013;57(1):40–7.

39. Kim M, Gans JD, Nogueira C, Wang A, Paik JH, Feng B, et al. Comparative oncogenomics identifies NEDD9 as a melanoma metastasis gene. Cell. 2006;125(7):1269–81.

40. Whittle JR, Wheatley AK, Wu L, Lingwood D, Kanekiyo M, Ma SS, et al. Flow cytometry reveals that H5N1 vaccination elicits cross-reactive stem-directed antibodies from multiple Ig heavy-chain lineages. J Virol. 2014;88(8):4047–57.

41. Wu X, Yang ZY, Li Y, Hogerkorp CM, Schief WR, Seaman MS, et al. Rational design of envelope identifies broadly neutralizing human monoclonal antibodies to HIV-1. Science. 2010;329(5993):856–61.

42. Brochet X, Lefranc MP, Giudicelli V. IMGT/V-QUEST: the highly customized and integrated system for IG and TR standardized V-J and V-D-J sequence analysis. Nucleic Acids Res. 2008;36(Web Server issue):W503–8.

43. Kearse M, Moir R, Wilson A, Stones-Havas S, Cheung M, Sturrock S, et al. Geneious Basic: an integrated and extendable desktop software platform for the organization and analysis of sequence data. Bioinformatics. 2012;28(12):1647–9.

44. High throughput HIV-1 microneutralization assay [Internet]. 2013. Available from: [+https://doi.org/10.1038/protex.2013.069+].

45. Montefiori DC. Measuring HIV neutralization in a luciferase reporter gene assay. Methods Mol Biol. 2009;485:395–405.

