## Supplemental Figures 1-6 for "Application of B cell immortalization for the isolation of antibodies and B cell clones from vaccine and infection settings"

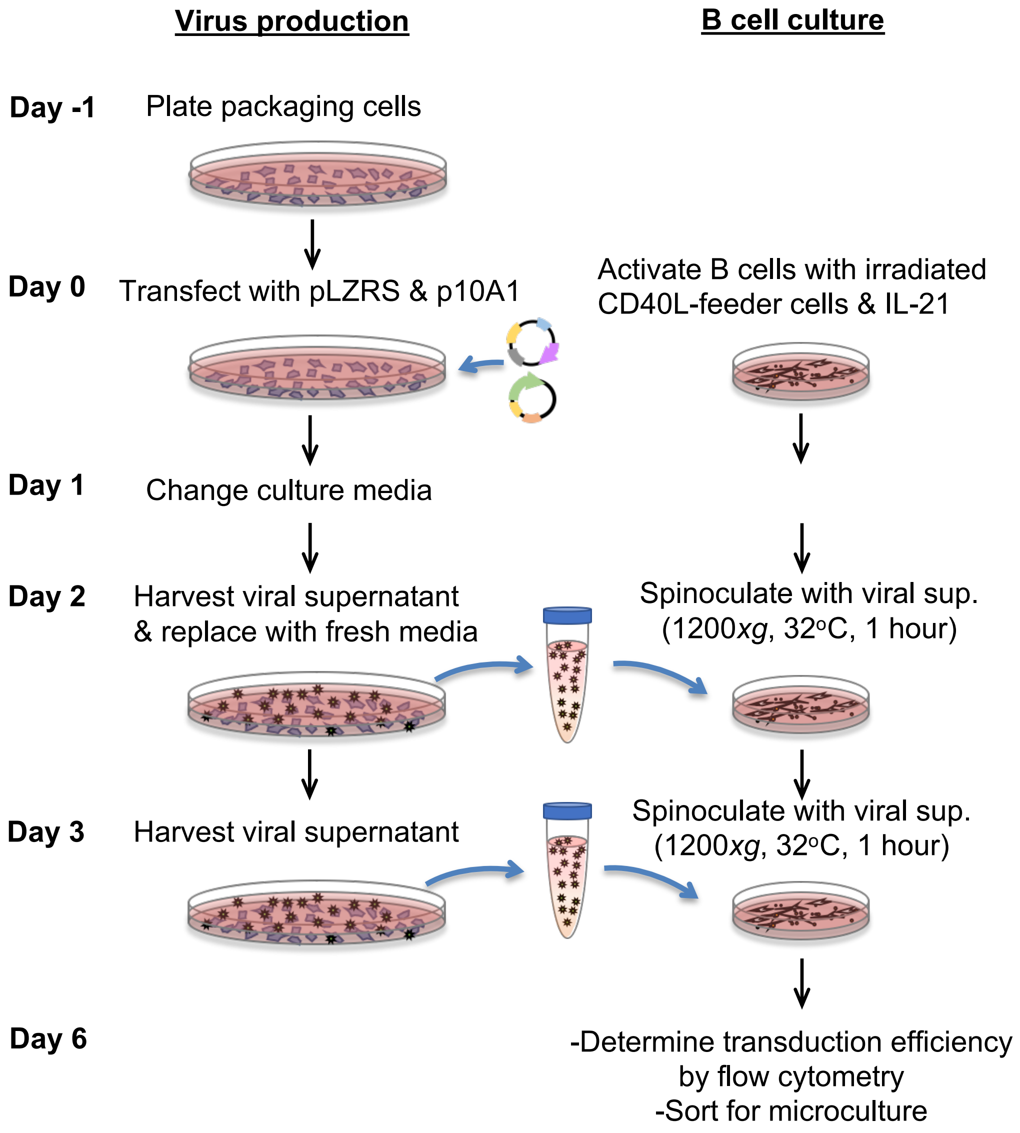


**Supplemental Figure 1. Schematic outlining the protocol for transducing primary B cells**

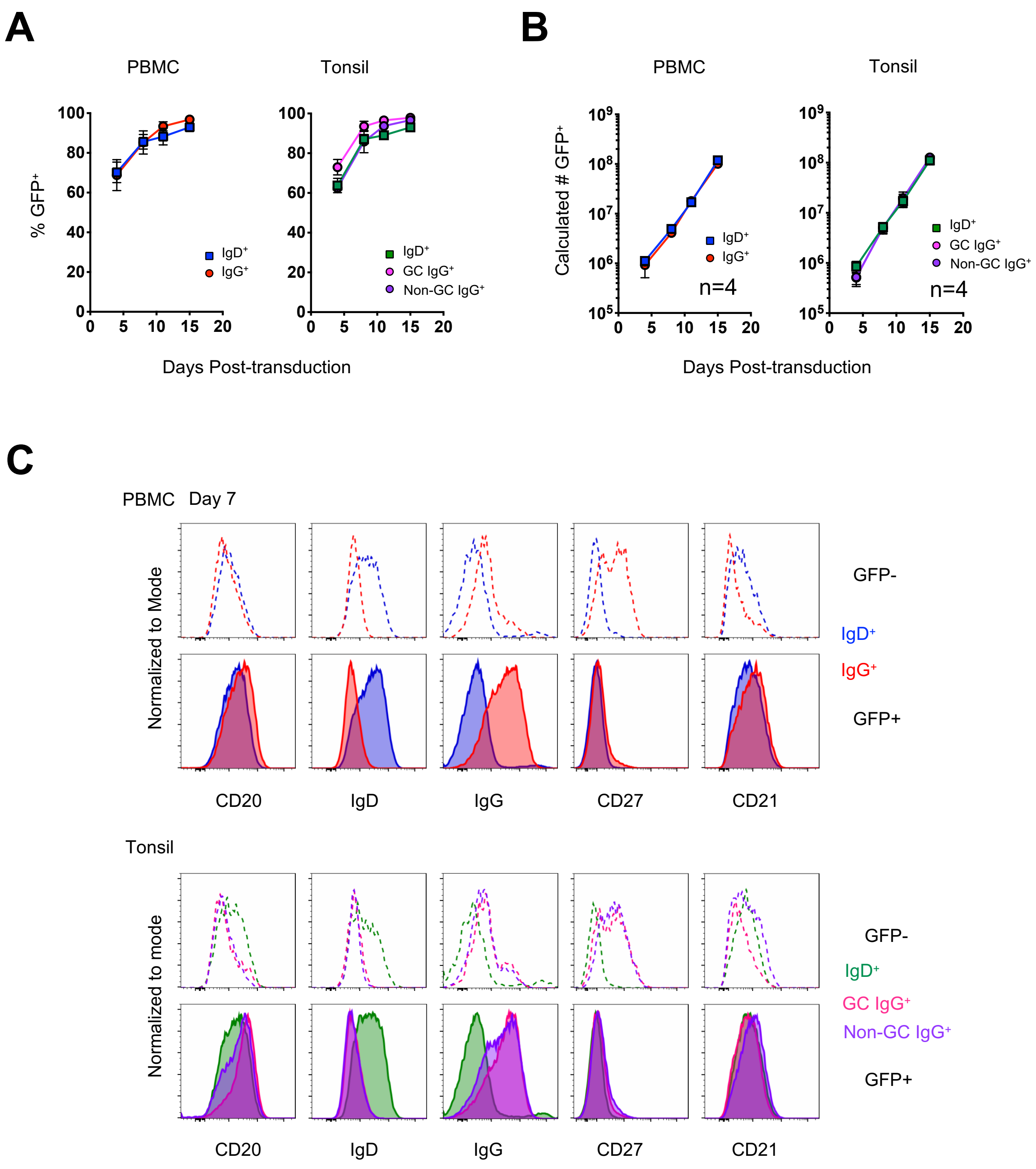


**Supplementary Figure 2. Transduced B cells expand and continue to express surface immunoglobulin.** (A) Following transduction, GFP⁺ B cells expand within the culture (PBMC: n=4; Tonsil: n=4). (B) GFP⁺ B cells were counted by flow cytometry and the total number of GFP⁺ B cells was calculated to account for splitting the cells 1:4 every 3 to 4 days. (C) Representative flow histograms of B cell cultures 7 days after transduction. B cells were gated based on expression of GFP and analyzed for surface expression of the indicated markers for each transduced B cell subset.

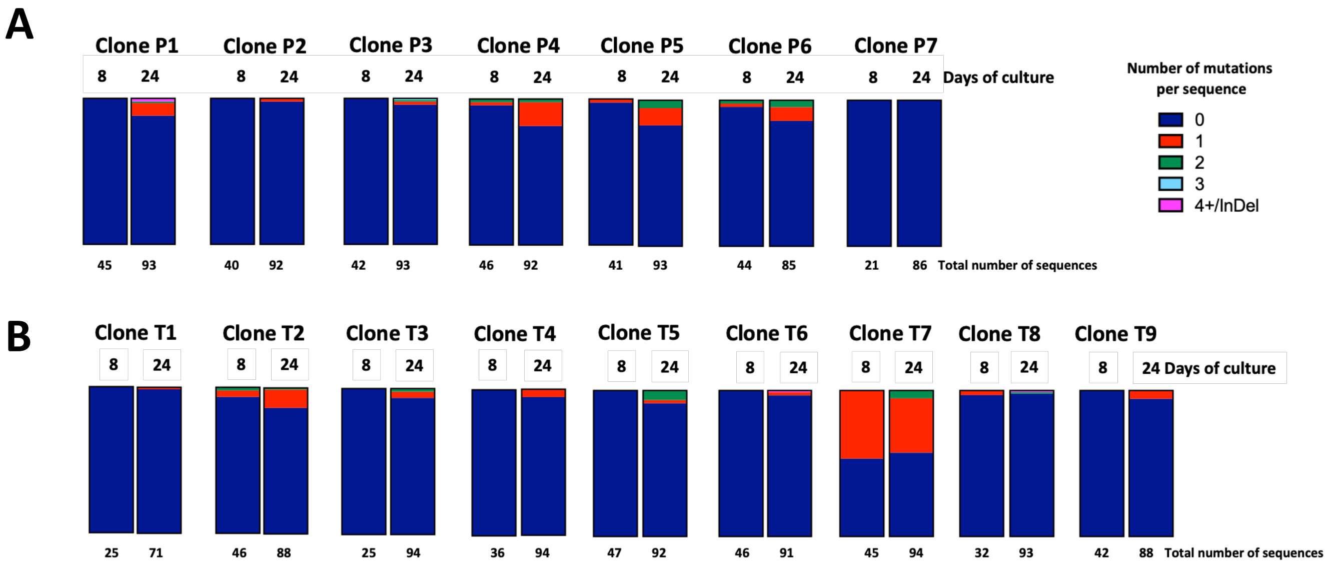


**Supplementary Figure 3. Acquisition of mutations within individual clones during *in vitro* culture.** (A, B) The frequency of individual sequences within a clonal population that contain zero, 1, 2, 3 or 4 or more mutations on day 8 and day 24 of culture for $IgG^+$ B cells from PBMC (A) and $IgG^+$ germinal center B cells from tonsil (B). The total number of sequences analyzed is indicated underneath.

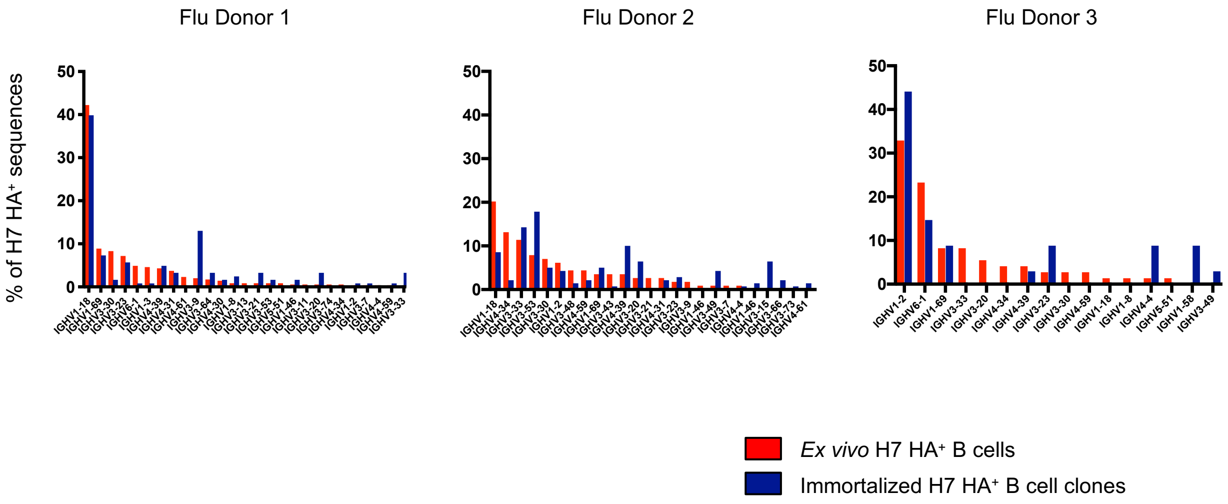


**Supplementary Figure 4. VH gene usage of H7 HA-specific B cells from *ex vivo* sorted B cells versus after B cell immortalization and clonal expansion.**

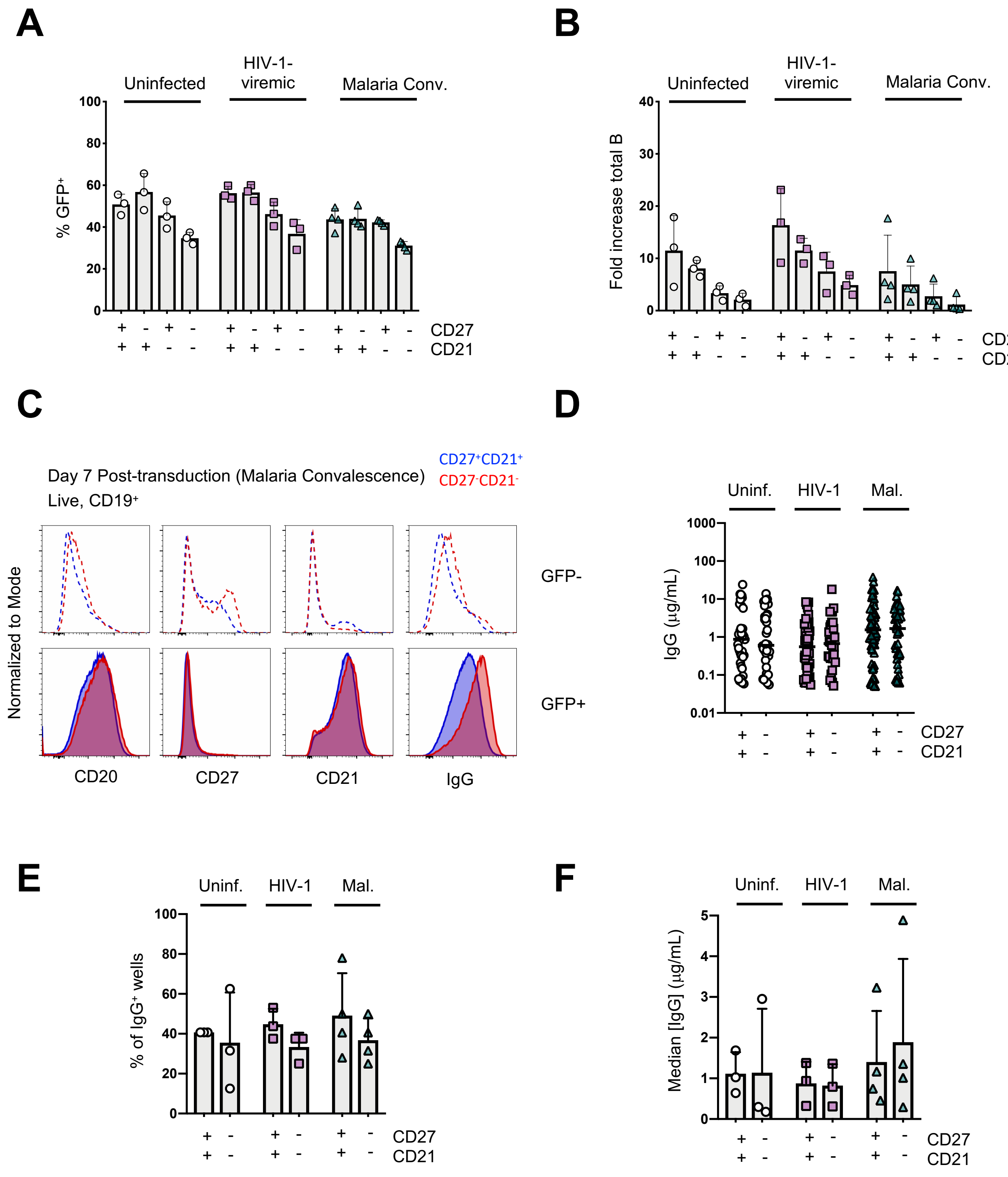


**Supplementary Figure 5. Immortalization of B cell subsets based on CD27 and CD21 expression from uninfected, HIV-1-viremic and following malaria infection.** (A) The frequency of GFP$^+$ B cells following transduction of the indicated subsets on day 4 post-transduction. (B) The fold-increase in B cell numbers during activation and transduction for the isolated B cell populations. (C) Representative flow histograms of B cell cultures 7 days after transduction. B cells were gated based on expression of GFP and analyzed for surface expression of the indicated markers, for the indicated B cell subsets. (D) Scatter plot showing the concentration of IgG in supernatants from IgG-positive wells on day 24 of microculture. (E) The frequency of IgG$^+$ wells after 24 days of microculture as determined by ELISA. (F) The median IgG concentration of IgG$^+$ wells on day 24 of microculture.

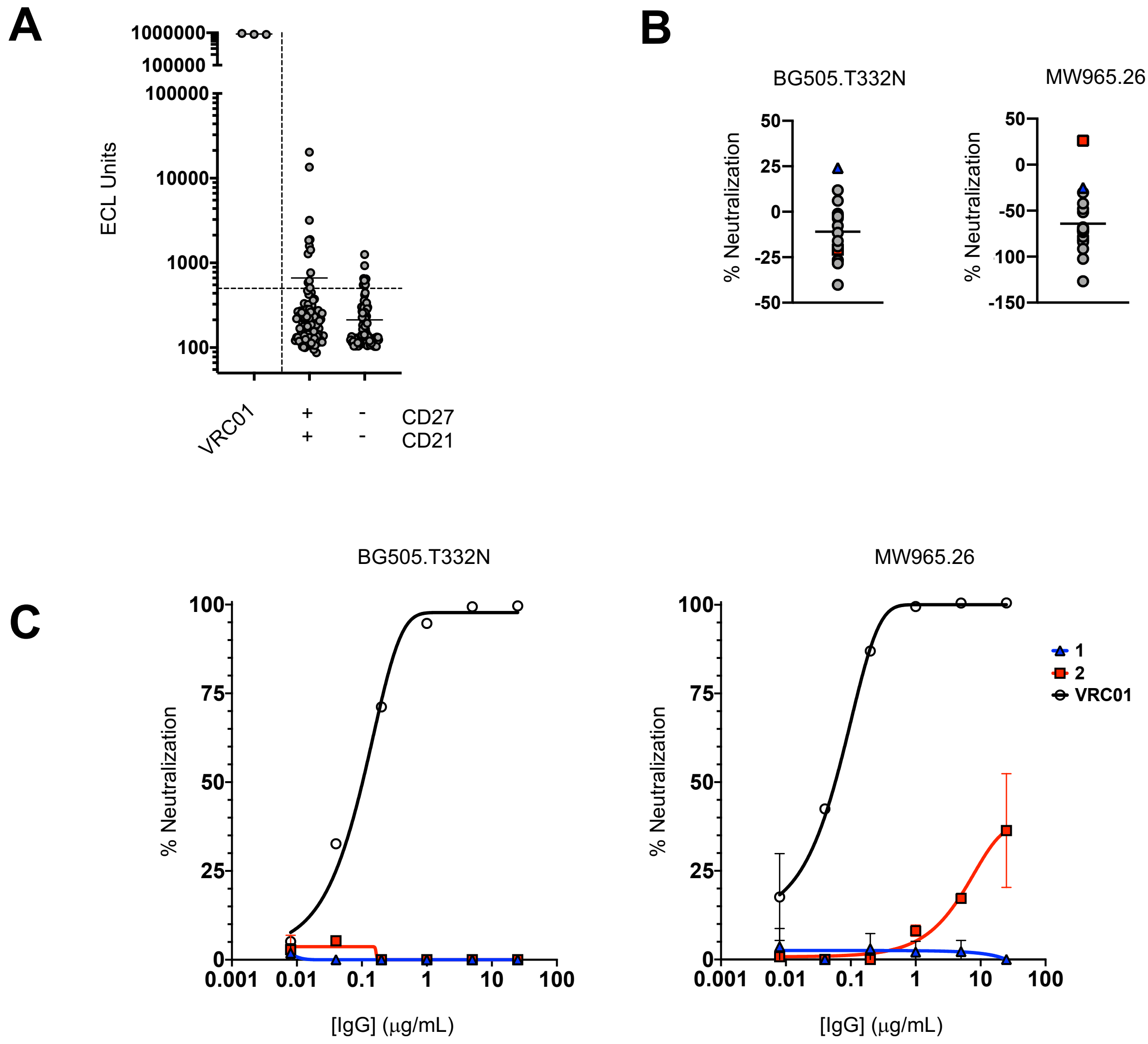


D

| Well | Donor | Viral Load (copies/mL) | Population | IGHV | % mutation |
|---|---|---|---|---|---|
| 1 | HIVpos1 | 385 | CD27⁺CD21⁺ | IGHV4-39 | 5.5 |
| 2 | HIVpos1 | 385 | CD27⁻CD21⁻ | IGHV1-24 | 4.2 |

**Supplemental Figure 6. Isolation of a HIV-1-specific antibody from the CD27$^-$CD21$^-$ B cell subset** (A) Supernatants from the CD27$^+$CD21$^+$ and CD27$^-$CD21$^-$ populations from HIV-1-viremic donors were screened for IgG binding to the BG505 SOSIP trimer. (B) Select wells were expanded based on their binding to the BG505 SOSIP trimer and supernatants from the expanded clones were screened for neutralizing activity against the MW965.26 and BG505 T332N pseudovirus strains in a microneutralization assay. (C) Wells 1 and 2 were further expanded, the immunoglobulin was purified and tested in the TZM-bl neutralization assay against the MW965.26 and BG505 T332N pseudovirus strains. (D) Immunogenetic characteristics of individual B cell sequences from B cells isolated from wells 1 and 2.
